# Monaural and binaural masking release with speech-like stimuli

**DOI:** 10.1101/2021.11.07.467581

**Authors:** Hyojin Kim, Viktorija Ratkute, Bastian Epp

## Abstract

The relevance of masking release by comodulation and interaural phase difference (IPD) for speech perception is still unclear. We used speech-like stimuli to link spectro-temporal properties of formants with masking release. The stimuli comprised a tone and three masker bands centered at formant frequencies F1, F2, and F3 derived from a consonant-vowel (/gu/, /fu/, and /pu/). The target was a diotic or dichotic frequency-modulated tone following F2 trajectories. Results showed no statistically significant comodulation masking release (CMR), while the binaural masking level difference (BMLD) was comparable to previous findings. The data suggest that factors other than comodulation may play a dominant role in grouping frequency components in speech.

## 1. Introduction

In a complex acoustic environment, frequency components of multiple sound sources are mixed in time. Nevertheless, the auditory system can separate and group frequency components into a target sound and a masking noise. The auditory system uses distinctive features of the target sound to form separate streams, such as “harmonicity”, “proximity” or “similarity” of its frequency components [Bregman, 1994, Bizley and Cohen, 2013, for a review]. When frequency components in stimuli are close to each other in frequency or time, they tend to be grouped into one stream. For example, when two pure tones are played close together in frequency or time, they are perceived as a single stream rather than two (e.g., trill threshold) [Miller and Heise, 1950]. However, the frequency components of natural sounds are often too far apart to be grouped solely by spectral proximity. Speech exhibits multiple prominent frequency components known as formants [Raphael et al., 2007]. The vowel /i/ can have three formants (F1, F2, and F3) at 270 Hz, 2290 Hz, and 3010 Hz, respectively [Raphael et al., 2007]. Whether these frequency components are grouped into a single stream by spectral proximity is unknown. Besides spectral proximity, formants also show coherent amplitude modulation patterns as a result of vocal tract vibration [Raphael et al., 2007]. This comodulation can serve as another cue for grouping by similarity for the auditory system, which may play a role in speech perception [Nelken et al., 1999]. Supporting the ecological relevance of comodulation, previous studies on the discrimination of speech sounds found that dyslexic children showed insensitivity to amplitude modulation rather than to formant changes [Ziegler and Goswami, 2005, Goswami et al., 2011]. Spatial continuity can be another entity for similarity that formants of one speech source share. For instance, speakers remain relatively constant in their spatial location [Bregman, 1994]. When continuous speech is segmented and presented alternatively to left and right ears, the integration of speech is hindered depending on the rate of speech alternation [Cherry and Taylor, 1954].

In psychoacoustical experiments, the benefits of both comodulation and spatial cues have been demonstrated as enhanced target detection performance. Regarding comodulation, the detection threshold of a tone masked by comodulated noise is lower compared to the threshold of a tone in noise with uncorrelated modulation patterns [Hall et al., 1984]. The difference in masked threshold is commonly referred to as comodulation masking release (CMR) [Hall et al., 1984]. Furthermore, adding comodulated noise components that are remote from the masked tone in frequency allows the detection threshold to be further decreased, resulting in increased CMR [Hall et al., 1990]. CMR paradigms can be divided into two main groups commonly referred to as “band-widening experiments” and “flanking-band experiments”. In the band-widening experiment, a single band of noise is used as a masker centered around the signal frequency. This masker is either modulated or uncorrelated. Masked thresholds are measured as a function of the masker bandwidth [Hall et al., 1984, Carlyon et al., 1989]. Masked threshold decreases with increased masker bandwidth, even when exceeding the critical bandwidth [Hall et al., 1984], indicating that CMR processing occurs across frequency channels. In the flanking-band paradigm, a masker consists of the signal-centered band and several narrow bands remotely centered around the target signal with spectral spacing (flanking bands, FBs). From the studies using the flanking-band paradigm, a general agreement is that CMR increases with increased spectral proximity between the target signal and FBs, and the number of FBs [Hall et al., 1990]. Both experiment paradigms showed the large benefit of comodulation in target detection and the effective frequency range of CMR processing. However, the frequency settings of CMR stimuli are very dissimilar to speech. Unlike the static frequency components in most CMR experiments, each phoneme in speech has specific formant settings in frequency and shows a formant transition from one phoneme to another. Few CMR studies used sweeping stimuli with a certain rate or frequency components with random-center frequency fluctuations [Verhey et al., 2013, Doleschal and Verhey, 2020]. Still, these stimuli are not similar to formant changes in speech. Similar to comodulation, interaural disparities in the masker or the signal can induce a masking release. The difference in masked threshold in the absence and presence of interaural disparities is referred to as binaural masking level difference (BMLD). For example, an interaural phase difference (IPD) of a pure tone in a narrow-band masker, this effect can be as high as 20 dB [e.g. van de Par and Kohlrausch, 1999]. The ecological validity of binaural cues has been shown by measuring speech intelligibility of masked speech signals in noise. Introduction of interaural differences leads to a difference in intelligibility of speech around 5 dB [Ho et al., 2015, Johansson and Arlinger, 2002]. This effect, measured at supra-threshold levels, is referred to as binaural intelligibility-level difference (BILD). With artificial stimuli, presenting both comodulation and interaural phase differences (IPD) showed a superposition of masking release at threshold [Epp and Verhey, 2009, Hall III et al., 2011]. However, little is known about how these two cues induce masking release with speech-like stimuli.

In this study, we aim to investigate the interplay of comodulation and IPD cues with the presence of speech-like formant transitions. We assumed that the fundamental unit of speech is a syllable rather than a phoneme [see e.g., Goswami et al., 2011, Vazeux et al., 2020]. We also assumed that masking release induced by a specific cue is an indicator of the presence of grouping, supporting the target separation from a masking noise. To design speech-like formant transitions, we analyzed speech samples of consonant and vowel combinations (CV). We extracted formant patterns and created three masker bands centered at formant frequencies F1, F2, and F3 based on CV combination: /gu/, /fu/, and /pu/. The generated masker bands had different degrees of center frequency transitions and level of covariation of center frequency: i) a stationary formant transition with minimal changes in center frequency (/gu/), ii) a gradual and covarying formant transition across the three masker bands (/fu/), and iii) a highly variable and independent formant transition across the three masker bands (/pu/). The target signal was a frequency modulated tone centered at F2, following the formant trajectory. To induce BMLD, we added an IPD of *π* to the target signal. Given that formant transitions, comodulated intensity fluctuations, and interaural differences are common properties of speech, we hypothesized that: a) spectro-temporal transitions in speech-like stimuli are beneficial for grouping, b) comodulation provides an additional grouping cue to spectro-temporal transitions (expressed as CMR), and c) binaural information in the form of IPD can be utilized as an additional beneficial cue on top of comodulation (expressed as BMLD), aligning with previous findings using a fixed flanking-band paradigm [e.g. Epp and Verhey, 2009].

## 2. Methods

### 2.1. Stimuli

We designed a masking experiment with stimuli that utilize formant patterns extracted from speech samples /gu/, /fu/, and /pu/ as a masker and target tone (Figure 1). For the masker, a slowly varying amplitude modulation was imposed on the spectral components. The target signal was a frequency modulated tone that followed the spectro-temporal transitions of the second formant, but was not amplitude modulated. Speech samples were extracted from the Oldenburg Logatome Corpus (OLLO) speech database [Wesker et al., 2005]. We chose VCV combination syllables with German speakers with no dialect. We chose one female speaker for analysis to preserve the nature of the speech signal. Due to the differences in formant transitions across speakers, averaging sound samples would not reflect natural formant transitions. We extracted CV combinations with a vowel detection algorithm to find the onset and endpoint of each phoneme [Garnaik et al., 2020]. To extract F1, F2, and F3 formants from the sound samples, the PRAAT software package was used [Boersma, 2011]. Five formants per frame were extracted within in the frequency range up to 5500 Hz. The duration of the analysis window length was 0.01 seconds. The first three formants were chosen as they are relevant for speech perception [Raphael et al., 2007]. Following the trajectories of the formants, we placed masker bands where each center frequency follows formant patterns (F1, F2, and F3) of /gu/, /fu/, and /pu/. The duration of the signal was 500 ms, including 20 ms raised-cosine ramps at the masker onset and offset.

**Figure 1:**
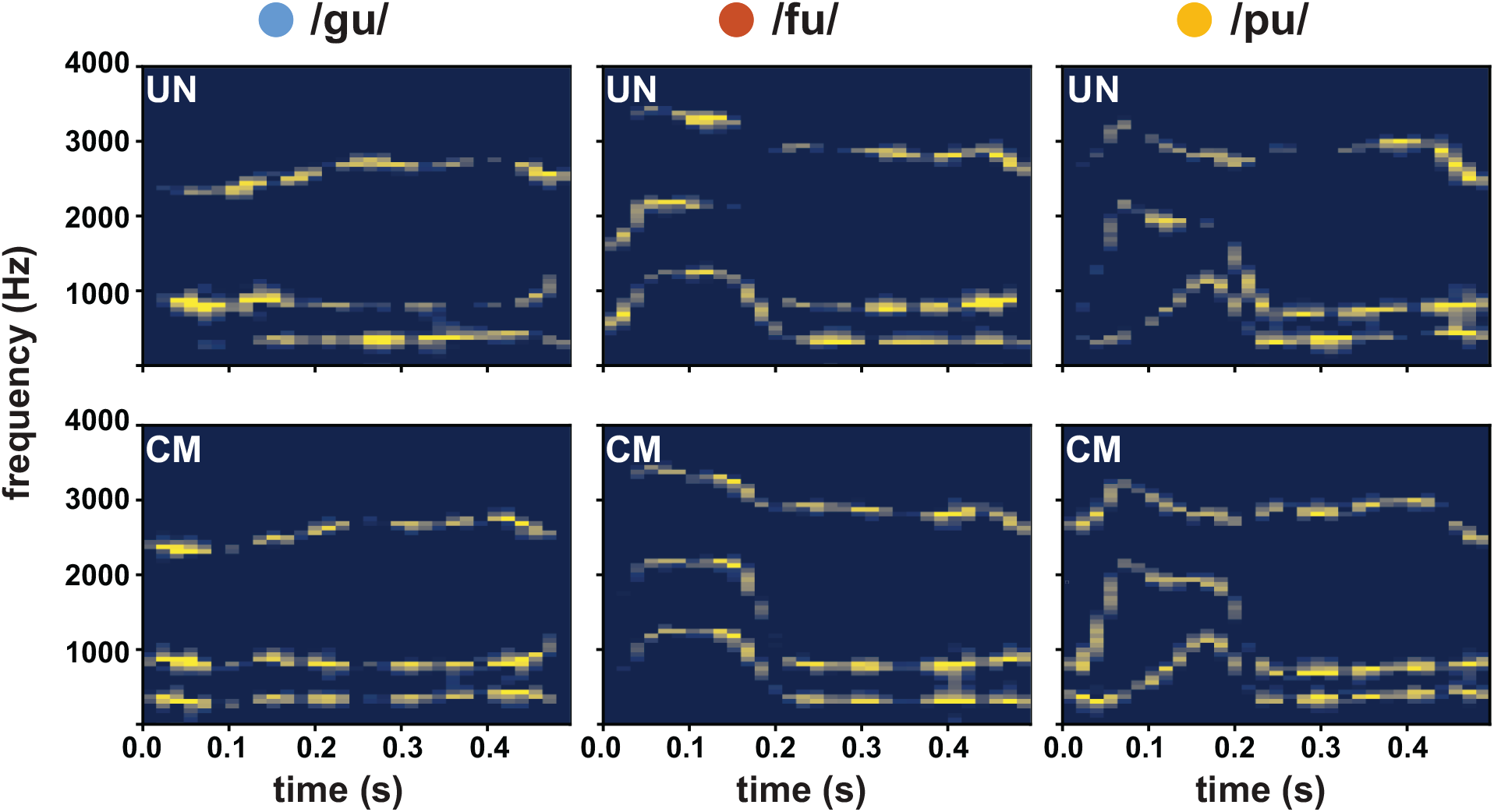
Spectrograms of the stimuli: /gu/, /fu/, and /pu/ from left to right. The top three panels show uncorrelated masker types, and the bottom three panels show comodulated masker types.

All the signals were generated digitally at a sampling rate of 44.1 kHz in MATLAB 2018b (TheMathworks, Natick, MA). The instantaneous frequency extracted from the formant analysis with PRAAT was used as a carrier and multiplied with a low-pass-filtered noise with a bandwidth of 20 Hz (without DC component) in order to generate narrow-band noise masker components *m*_*i*_ with slow amplitude modulations:

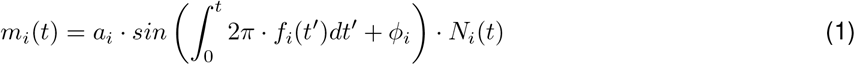

where *N*_*i*_ is low-pass-filtered noise without a DC component. *a*_*i*_ represents the amplitude of one masker band, scaled such that the overall level of the masker was 60 dB SPL. *ϕ*_*i*_ represents the starting phase of the carrier, chosen randomly from a normal distribution for each band. *f*_*i*_ represents the time-varying frequency estimated from the formant analysis. For the uncorrelated (UN) noise, a different modulator was generated and multiplied with each time-varying frequency band (formant) separately. For the comodulated (CM) noise, the same modulator was used for all masker bands. The target tone was added, which followed the center frequency of the middle masker band:

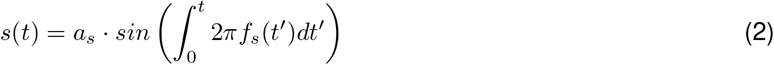

where *a*_*s*_ is the amplitude and *f*_*s*_ is the signal frequency at each time point.

### 2.2. Protocol

After the training session (one presentation of each condition), each listener performed three threshold measurements for all conditions. For each measurement, conditions were presented in randomized order. The mean thresholds of each condition were calculated as the average of three track estimates. One additional measurement was conducted if the thresholds from the last three measurements had a high standard deviation (SD *>* 3 dB). During the threshold measurement, listeners were seated in a double-walled, soundproof booth with ER-2 headphones. We used an adaptive, three-interval, three-alternative forced-choice procedure (3-AFC) implemented in MATLAB 2018b (The Mathworks, Natick, MA) with a one-up, two-down rule to estimate the 70.7% of the psychometric function [Ewert, 2013, Levitt, 1971]. Three sound intervals were presented with a pause of 500 ms in between. Two intervals contained the maskers only. The remaining interval contained the target tone with maskers. The listeners’ task was to choose the interval with the target tone by pressing the corresponding number key (1, 2, 3) on the keyboard. Visual feedback was provided, indicating whether the answer was “WRONG” or “CORRECT.” The target tone’s start level was set to 75 dB SPL, and the tone level was adjusted with the initial step size of 8 dB. The step size was halved after each lower reversal until it reached the minimum step size of 1 dB. The mean of six reversals with the minimum step size was estimated as the threshold.

### 2.3. Listeners

From hearing screening, we recruited ten normal-hearing listeners. None of them reported any history of hearing impairment and had pure-tone hearing thresholds within 15 dB HL for the standard audiometric frequencies from 125 to 4000 Hz.

## 3. Results

Masked thresholds for all combinations of parameters are shown in Figure 2.

**Figure 2:**
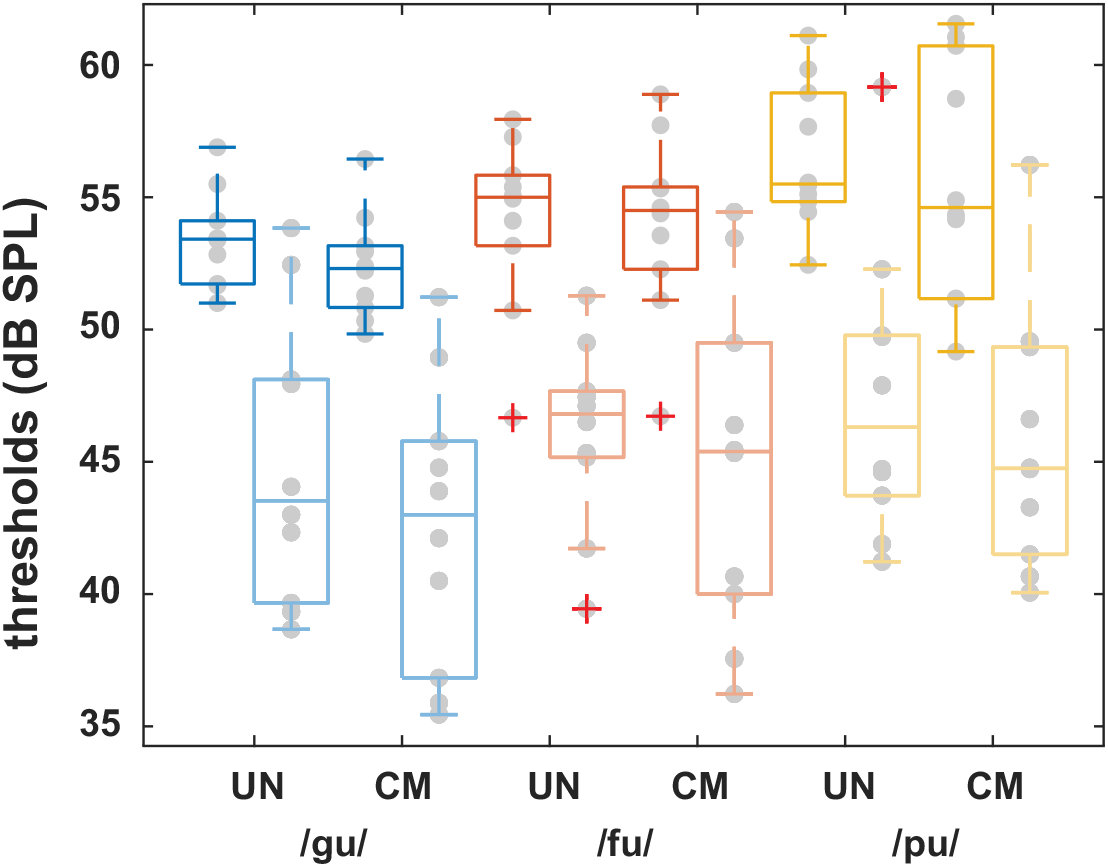
Individual and mean masked thresholds for three masker types: /gu/ (blue), /pu/ (orange), and /fu/ (yellow). Each masker type was either uncorrelated (UN) or correlated (CM). Additionally, the target tone was presented with IPD of 0 or *π*. The thresholds of conditions with IPD of *π* are plotted in lighter color. Individual mean masked thresholds are represented with grey points. Crosses indicate outliers in the data.

A generalized linear model (GLM) analysis revealed no significant effect of modulation type (*β* = −0.66, *SE* = 1.08, *p* = 0.54). The binaural cue (*β* = −0.04, *SE* = 0.013, *p <* 0.05) and the CV type (*β* = 1.52, *SE* = 0.47, *p <* 0.05) showed significant effects on thresholds. No significant interaction was found between modulation type and binaural cue (*β* = −0.33, *SE* = 0.14, *p* = 0.022).

Based on the thresholds, we calculated CMR and BMLD for each of the speech-like stimuli. Results are shown in Figure 3. The mean CMR for each condition was: /gu/ (1.03 dB), /fu/ (0.84 dB), /pu/ (0.11 dB) in diotic the conditions, and /gu/ (2.4 dB), /fu/ (1.83 dB), and /pu/ (1.22 dB) in the dichotic conditions. For /gu/, CMR in both diotic and dichotic conditions were significantly different from zero (t-test, *p <* 0.05). For /fu/, the CMR was not significantly different from zero in any of the conditions. For /pu/, only the CMR in the dichotic condition was significant from zero (t-test, *p <* 0.05). No significant difference was found between all conditions (one-way ANOVA, in diotic condition: *F* (5, 54) = 0.82, *p >* 0.05). CMR values showed a high variability in /fu/ and /pu/ maskers (Figure 3a with grey scatter plot). For /fu/ with IPD of 0, CMR values ranged from - 6.11 dB to 5.28 dB. For /pu/ with IPD of *π*, CMR values showed high variability ranging from - 6.8 dB to 10.3 dB (individual data are available in the Supplement). The mean BMLD for each condition was: /gu/ (8.46 dB), /fu/ (8 dB), /pu/ (9.83 dB) with UN maskers, and /gu/ (9.01 dB), /fu/ (10.02 dB), and /pu/ (9.04 dB) with CM maskers (Figure 3b). No significant difference was found across all CV combinations in both diotic and dichotic conditions (one way ANOVA, *F* (5, 54) = 0.25, *p >* 0.05).

**Figure 3:**
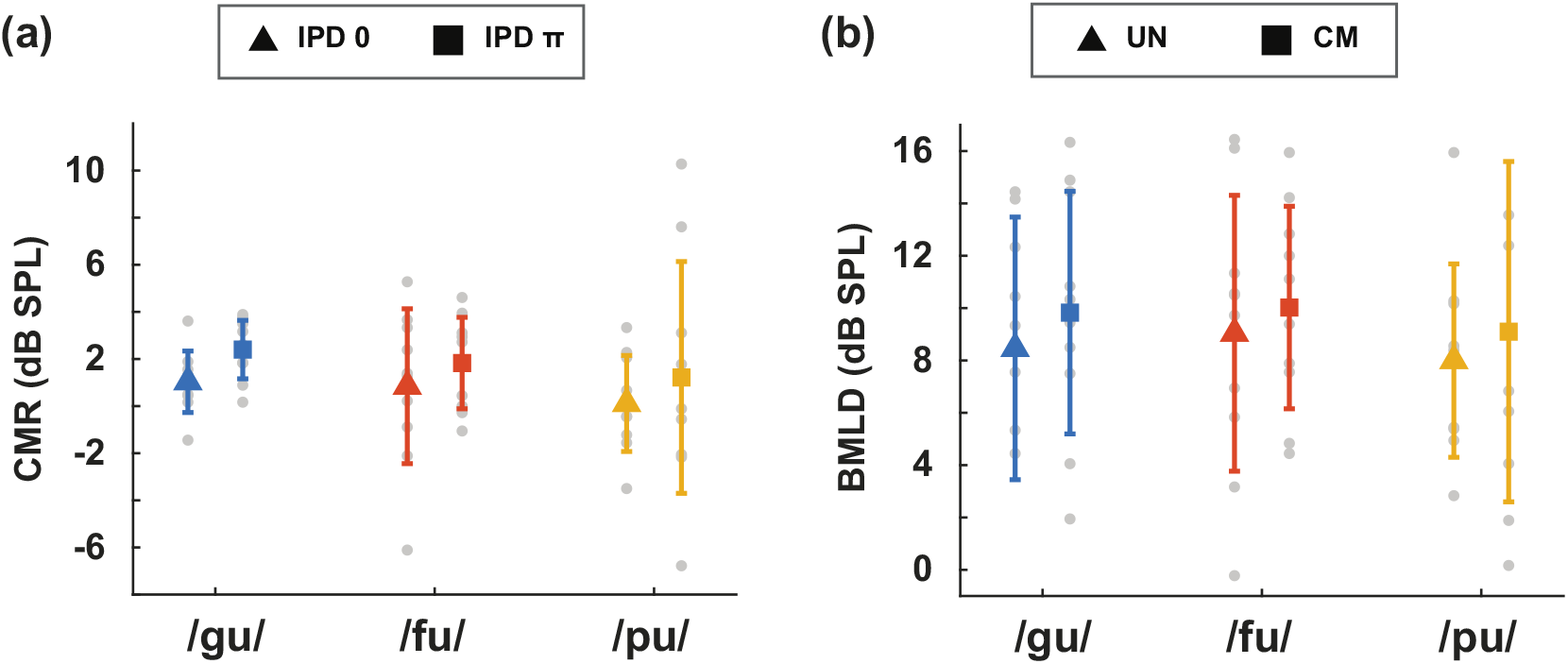
CMR and BMLD for all conditions. (a) The mean CMR for each condition. Triangles indicates CMR with an IPD of 0, squares indicates CMR with an IPD of *π*. (b) The mean BMLD for each condition. Triangles indicates BMLD for UN masker type, squares indicates BMLD for CM masker type. Blue color represents /gu/, orange color represents /fu/, and yellow color represents /pu/. Error bars indicate plus-minus one standard deviation.

Thresholds, CMR, and BMLD showed high variability across listeners. Thresholds in diotic conditions showed less variability than thresholds in dichotic conditions (see Figure 2), and introduction of comodulation increased variability of thresholds for the /pu/ condition. Hence, variabilites in thresholds and differences in the effect of IPD and comodulation underlie the variability in the derived measures of CMR and BMLD.

## 4. Discussion

In this study, we investigated the ecological relevance of masking release, specifically CMR and BMLD. For BMLD, the results were in line with data on voiced vowels where BMLD was in the range from 7.3 to 11.5 dB [Ho et al., 2015]. The results were also consistent with binaural intelligibility level difference (BILD) between 5.7 and 7.7 dB using speech signals [Johansson and Arlinger, 2002]. However, CMR values were not significantly different to zero, in contrast to previous CMR studies with non-stationary stimuli. In a previous study by Verhey et al. [2013], they measured CMR with sweeping maskers. Proportional to the increase in sweep rate, the CMR was reduced (e.g., from 11 dB to 6 dB). In another CMR study with random-variations in frequencies of maskers, CMR was small (e.g., 5 dB) compared to a CMR study with stationary stimuli (e.g., 6 – 10 dB) [Doleschal and Verhey, 2020]. The stimuli used in this experiment had different degrees of formant variation (stationary, correlated formant transitions, uncorrelated formant transitions). For the condition with stationary formant changes (e.g., /gu/), the individual CMR was typically less than 3 dB while CMR with more formant variations (e.g., /fu/, /pu/) showed individual values ranging from - 6 dB to 10 dB.

Based on previous studies by Hall et al. [1990], the number of FBs may play a role in facilitating the grouping of formants by comodulation. One of the differences in the stimulus design of the current study compared to previous studies was the number and the spectral proximity between flanking bands. While the study by Verhey et al. [2013] with sweeping masker had five masker bands, the stimuli used in this study had three FBs and a wider spacing than the sweeping masker. This factor may be related to the reduced amount of CMR in our study. Another difference in stimuli was non-stationary frequency changes with formant patterns (e.g., /fu/ and /pu/). While previous studies used either randomly varied frequency in time or the same sweeping patterns across noise bands, we extracted formant changes from speech samples. With more formant variations, some listeners showed higher positive CMR or negative CMR than the stimuli with stationary formant patterns (e.g., /gu/). In the condition with /fu/ and IPD of 0, the maximum value of CMR was 5.28 dB while the minimum value of CMR was -6.11 dB. Similarly, in the condition with /pu/ and IPD of *π*, the maximum value of CMR was 10.28 dB, while the minimum value of CMR was -6.78 dB. This variability might indicate individual challenges in being able to utilize the cues.

The introduction of comodulation in /pu/ increased the standard deviation of the masked thresholds relative to UN /pu/. Condition /pu/ was the condition with the highest degree of variability in the formant transitions (see Figure 1). This is line with the idea that the high degree of variability might have challenged some listeners in the use of comodulation for masking release. Evaluating CMR and BMLD for /pu/ (Figure 3), the higher standard deviations in the dichotic conditions compared to the diotic conditions (CMR) and comodulated compared to the unmodulated conditions (BMLD) indicate that some listeners in this study could not utilize both comodulation and IPD to improve masked thresholds. Thus, the underlying neural basis for CMR and BMLD interaction needs to be further investigated to explain how formant transitions affect the use of IPD cues differently in /pu/ and /fu/ conditions.

## 5. Conclusion

In this study, we evaluated the ecological relevance of CMR and BMLD. Our hypothesis was that if CMR and BMLD are relevant to speech perception, speech-like stimuli will induce a comparable amount of CMR and BMLD to previous findings. Results showed that CMR was not significant while BMLD was around 9 dB. We concluded that comodulation was either not a sufficient cue in the present study to group three formants, or irrelevant for the task. We suggest that other sound properties such as formant transition patterns, the proximity of masker bands to the target signal, and the number of masker bands can be more relevant for grouping frequency components to induce speech perception.

